# Dysregulated CD38 expression in blood and skin immune cells of patients with hidradenitis suppurativa

**DOI:** 10.1101/2023.01.27.525867

**Authors:** Peter Dimitrion, Iltefat Hamzavi, Congcong Yin, Ian Loveless, Jugmohit Toor, Kalpana Subedi, Namir Khalasawi, Angela Miller, Richard Huggins, Indra Adrianto, Jesse Veenstra, Gautham Vellaichamy, Aakash Hans, Steven Daveluy, Mohammad Athar, Wilson Liao, Henry Lim, David Ozog, Li Zhou, Qing-Sheng Mi

**Affiliations:** Center for Cutaneous Biology and Immunology Research, Department of Dermatology, Henry Ford Health System, Detroit, MI, 48202, USA; Immunology Research Program, Henry Ford Cancer Institute, Henry Ford Health System, Detroit, MI, 48202, USA; Cancer Biology Graduate Program, School of Medicine, Wayne State University, Detroit, MI, 48202, USA; Center for Bioinformatics, Department of Public Health Sciences, Henry Ford Health System, Detroit, MI, 48202, USA; Department of Dermatology, Wayne State University School of Medicine, Detroit, MI, 48202, USA; University of Alabama at Birmingham (UAB) Research Center of Excellence in Arsenicals, Department of Dermatology, University of Alabama at Birmingham, Birmingham, AL, United States; Department of Dermatology, University of California San Francisco, San Francisco, California, USA; Department of Biochemistry, Microbiology, and Immunology, School of Medicine, Wayne State University, Detroit, MI, 48202, USA

**Keywords:** Hidradenitis suppurativa, Cytometry by time-of-flight, Monocytes, CD38, Natural killer cells, inflammation, Dendritic cells, imaging mass cytometry, endothelial cells

## Abstract

2.

**Background:** Hidradenitis suppurativa (HS) is a multifactorial, inflammatory skin disease. Increased systemic inflammatory comorbidities and serum cytokines highlight systemic inflammation as a feature of HS. However, the specific immune cell subsets contributing to systemic and cutaneous inflammation have not been resolved.

**Objective:** Identify features of peripheral and cutaneous immune dysregulation.

**Methods:** Here, we generated whole-blood immunomes by mass cytometry. We performed a meta-analysis of RNA-seq data, immunohistochemistry, and imaging mass cytometry to characterize the immunological landscape of skin lesions and perilesions from patients with HS.

**Results:** Blood from patients with HS exhibited lower frequencies of natural killer cells, dendritic cells, and classical (CD14+CD16-) and nonclassical (CD14-CD16+) monocytes, as well as higher frequencies of Th17 cells and intermediate (CD14+CD16+) monocytes than blood from healthy controls. Classical and intermediate monocytes from patients with HS had increased expression of skin-homing chemokine receptors. Furthermore, we identified a CD38+ intermediate monocyte subpopulation that was more abundant in the immunome of blood from patients with HS. Meta-analysis of RNA-seq data found higher CD38 expression in lesional HS skin than in perilesional skin, and markers of classical monocyte infiltration. Imaging mass cytometry showed that CD38+ classical monocytes and CD38+ monocyte-derived macrophages were more abundant in lesional HS skin.

**Conclusion:** Overall, we report targeting CD38 may be worth pursuing in clinical trials.

**Key Messages:**

1. Monocyte subsets express markers of activation in circulation and HS lesions
2. Targeting CD38 may be a viable strategy for treating systemic and cutaneous inflammation in patients with HS

**Capsule Summary:** Dysregulated immune cells in patients with HS express CD38 and may be targeting by anti-CD38 immunotherapy.

## Background

Hidradenitis suppurativa (HS) is a chronic inflammatory skin disease in which patients develop painful lesions that greatly impact quality of life^1^. HS prevalence is estimated to be about 1%, and trends in genetic ancestry and family histories point to a key role for inheritance^2^. Genetic studies have identified several disease-susceptibility loci^2^; however, no specific genes have been identified to explain the exuberant inflammatory response in patients with HS^2, 3^.

Although HS pathogenesis remains enigmatic, it is thought that subclinical inflammation promotes changes in cutaneous architecture and the immune compartment, leading to occlusion of hair follicles and formation of abscesses, which sets the stage for disease^3–9^. The transition from subclinical to clinically severe disease is thought to involve an inciting event (e.g., abscess rupture) that overwhelms the cutaneous immune system with immunogens, such as keratin debris and pattern/damage associated molecular patterns^9^. As acute inflammation is resolved, patients progress to a chronic inflammatory state, where the initial infiltrate is cleared and immune-stromal interactions reshape the cutaneous microenvironment^9^.

The number of different immune pathways and effector molecules implicated in HS has grown substantially^4, 10, 11^. Despite this, very few reliable biomarkers are known for accurately staging, prognosticating, and monitoring HS disease, and our understanding of how the dysregulated immune response is coordinated by effector molecules remains poorly understood^12^. Along with an increased prevalence of systemic inflammatory comorbidities, patients with HS also have higher levels of circulating inflammatory proteins^3, 5, 8^, and these findings highlight that systemic inflammation is a feature of HS disease. A dysregulated serum proteome may also affect the composition of the peripheral immunome and the activation status of circulating immune cells. Circulating immune cells serve as a good proxy for disease severity and risk of systemic comorbidities in other inflammatory conditions^3, 13, 14^. However, an unbiased characterization of the changes in the peripheral immunome of patients with HS has not been published. We hypothesized that patients with HS would harbor significantly perturbed peripheral immunomes and that distinct cell types or subsets may contribute to the systemic and cutaneous inflammation seen in these patients.

For studying complex inflammatory diseases, such as HS, that have multiple axes of inflammation and multifactorial etiology, bulk analyses are fruitful for identifying tissue-wide perturbations but are insufficient for systematically identifying cell-type specific changes that may underly pathogenesis. To date, some groups have employed single-cell sequencing and identified cell-specific changes in HS skin^10, 15, 16^, but no studies at the single cell level have been conducted on the peripheral immune system in HS.

In this study, we used mass cytometry by time-of-flight (CyTOF) to characterize peripheral immune cell changes in fresh blood samples from patients with HS. We found that patients with HS had dramatically decreased frequencies of natural killer (NK) cells, dendritic cells (DCs), and classical (CD14+CD16-; C.monos) and nonclassical (CD14-CD16+; NC.monos) monocytes, as well as significantly increased frequencies of Th17 cells and intermediate (CD14+CD16+; I.monos) monocytes. Patients with Hurley Stage II HS had markedly elevated CD38^+^ I.monos in circulation, which was the only cell population elevated in this group. CD38 expression was significantly higher in lesional HS skin than in perilesional skin, and the genes that were significantly correlated with CD38 expression were enriched for those expressed by C.monos and NK cells. Utilizing imaging mass cytometry (IMC), we found that endothelial cells (ECs), monocytes, and monocyte-derived macrophages (mono-macs) from lesional skin expressed higher CD38 than cells from perilesional skin. Monocytes and mono-macs colocalized to EC microenvironments that were functionally distinct from perilesional EC microenvironments. Collectively, our data uncover previously unreported peripheral and cutaneous immune dysregulation in patients with HS.

## Results

### Mass cytometry reveals novel dysregulation of circulating immune cell subsets

CyTOF studies of the peripheral immunome have uncovered disease-associated immune cell signatures and immunological drivers of disease^17, 18^. We used a standardized CyTOF assay to assess fresh whole blood from 18 patients with HS (6 Hurley Stage II; 12 Hurley Stage III) and 11 healthy control patients (HC) (**Table S1; Figure 1A**). T-distributed stochastic neighbor embedding (t-SNE) dimensionality reduction defined 11 major clusters of immune cells in whole blood, which were corroborated by lineage marker expression (**Figure 1B-C; Table S2**). Principal component analysis (PCA) showed minor separation between the HS and HC immune populations (**Figure S1A**).

**Fig.1.**
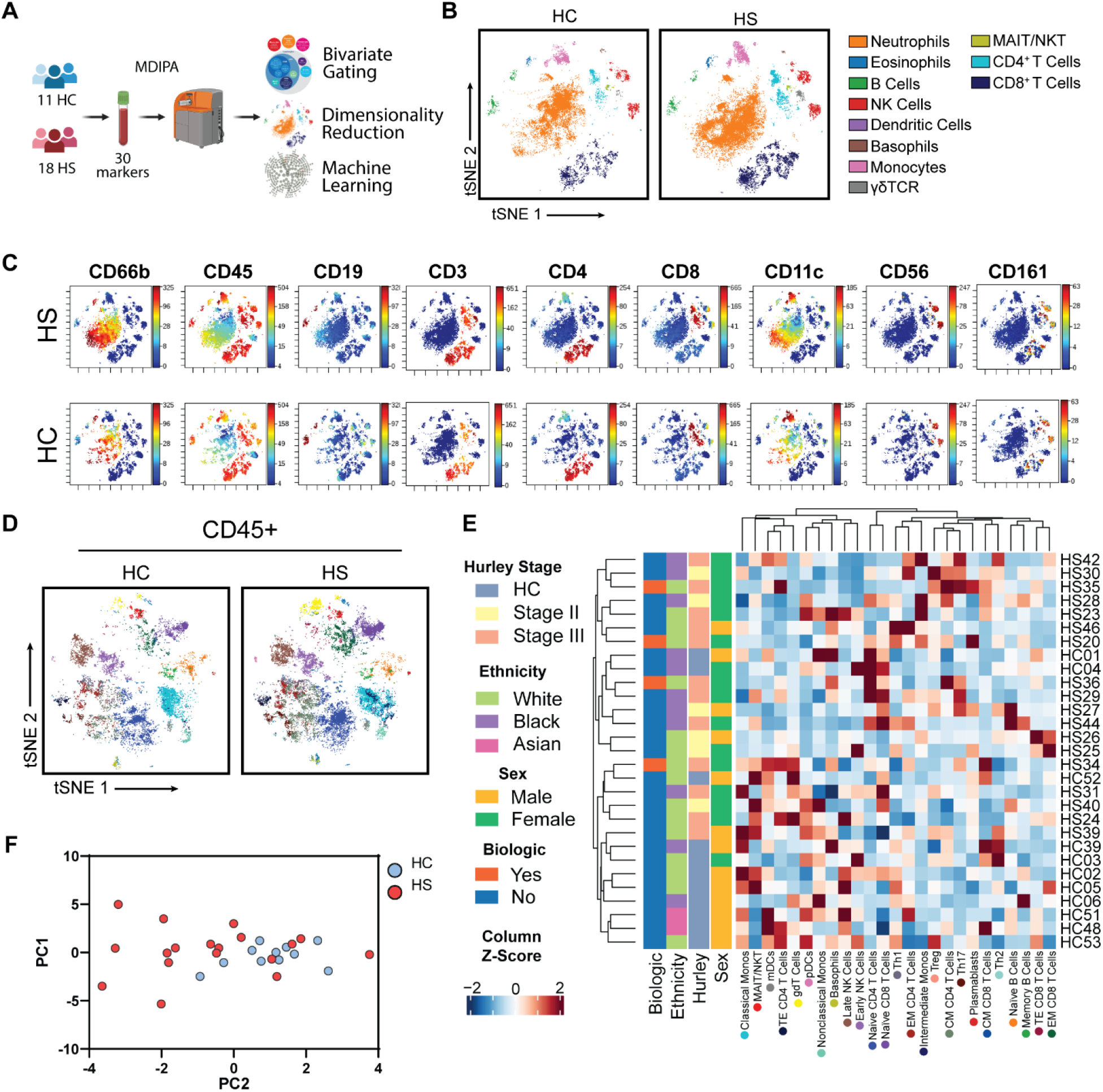
Characterization of the peripheral immunome of patients with hidradenitis suppurativa. (A) Schematic of CyTOF immune profiling experimental design; (B) Representative t-SNE-clustering of live singlets highlighted by major immune cell populations identified by bivariate gating; (C) Lineage marker expression on t-SNE plots; (D) Representative t-SNE-clustering of CD45^+^ cells highlighted by subset identity identified by bivariate gating; (E) Heatmap of Z-score normalized frequencies with hierarchical clustering. Colored dots below subset name correspond tSNE associated in Figure 1D; (F) Principle Components Analysis (PCA) plot of CD45^+^ subsets colored by HS and HC. Abbreviations: HC, healthy control; HS, hidradenitis suppurativa.

Neutrophils are thought to play a key role in HS pathogenesis^11, 19^. Interestingly, we found similar levels of circulating neutrophils in HS and HC samples (**Figure S1B**). Given that neutrophils constitute a large fraction of total blood leukocytes (40%-60%), we excluded them and performed deep immune profiling of CD45^+^ immune cell subsets.

Using bivariate gating, we identified 25 CD45^+^ cell subsets (**Figure 1D-1E; S2; Table S2)**. Hierarchical clustering of the frequencies of these subsets showed heterogeneity between the immunomes of patients with HS unexplained by use of biologic drugs, sex, ethnicity, or Hurley Stage (**Figure 1E**). PCA analysis of CD45^+^ immune subsets showed a more pronounced separation between HS and HC cell populations (**Figure 1F**).

We sought to define potential perturbations in the HS immunome. Among lymphoid-lineage cells, NK cells have been overlooked in studies of HS^9, 20^. Interestingly, we found a significant reduction in the frequency of circulating NK cells in HS (**Figure 2A**). And analysis of early and late NK cell subsets showed a decreasing trend in both early and late NK cells in HS (**Figure 2A**).

**Fig.2.**
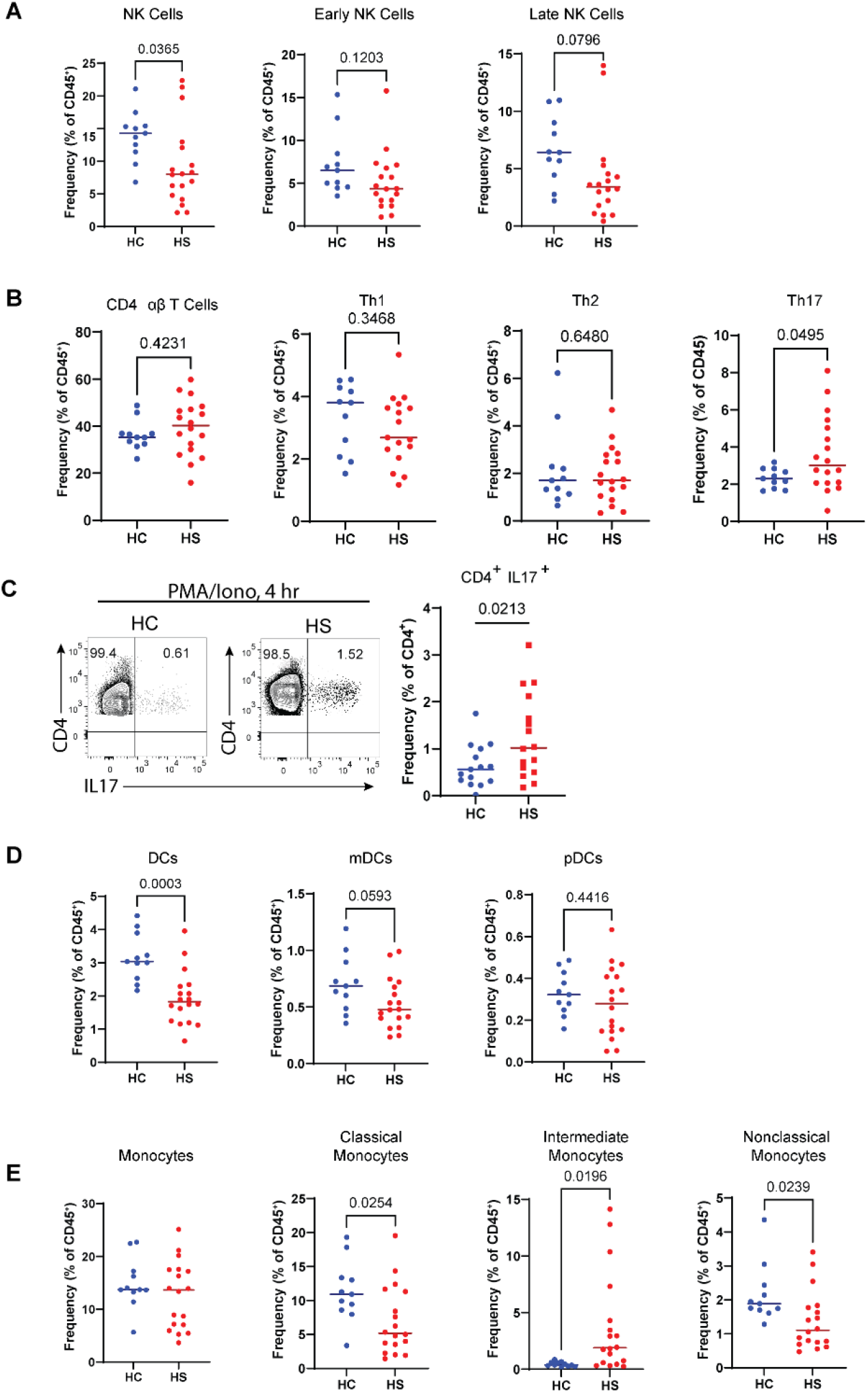
Dysregulated peripheral myeloid and lymphoid cell populations in patients with hidradenitis suppurativa: (A) Frequency of total NK, early NK, and late NK cells as a percent of CD45^+^ cells; (B) Frequency of total αβ T cells, Th1, Th2, and Th17 cells as a percent of CD45^+^ cells; (C) Representative bivariate flow cytometry plots of stimulated PBMCs (4 hours PMA/Iono) from HC and HS pre-gated on live singlet CD3^+^ CD4^+^ cells showing the expression of IL-17; (D) Frequency of total DCs, mDCs, and pDCs as a percent of CD45^+^ cells; (E) Frequency of total, classical, intermediate, and nonclassical monocytes as a percent of CD45^+^ cells. *P* values reported are the result of an unpaired students *t*-test. Abbreviations: DC, dendritic cells; NK, natural killer; HC, healthy control; HS, hidradenitis suppurative; mDC, myeloid dendritic cells; PBMC, peripheral blood mononuclear cells; pDC, plasmacytoid dendritic cells; PMA/Iono, phorbol myristate acetate and ionomycin.

No differences were observed in the frequency of total CD4^+^ αβT cells between HS and HC, but the frequency of the Th17 subset was elevated in HS (**Figure 2B**), which is consistent with previous reports implicating aberrant IL-17 mediated immunity in HS^3, 4, 8, 20, 21^. We then tested whether the functional capacity of Th17 cells would also be different between HS and HC by stimulating peripheral blood mononuclear cells (PBMCs) (from an independent cohort of patients with HS) with phorbol myristate acetate and ionomycin (PMA/Iono) for 4 hours. We found that patients with HS had a significantly greater frequency of CD4^+^ IL-17^+^ cells, indicating that circulating Th17 cells in patients with HS have an increased intrinsic ability to produce IL-17 (**Table S2; Figure 2C**). These results show that patients with HS have increased frequency and function of circulating Th17 cells.

Myeloid cells, such as monocytes and DCs, are recruited to sights of inflammation where they orchestrate the immune response at sites of tissue damage ^22^. In the myeloid compartment, we found that the frequency of total DCs was lower in HS (**Figure 2D**). Furthermore, the decreased levels of myeloid-DCs (mDCs) in HS approached significance (*p*=0.0593), whereas plasmacytoid-DCs (pDCs) showed no such trend (*p*=0.4416) (**Figure 2D**).

Monocytes serve many roles in the initiation and resolution of inflammation, tissue repair, and they are progenitors of antigen presenting cells in the skin ^22–25^. We found no difference in the total monocyte frequency in the blood of patients with HS relative to HC (**Figure 2E**). Further analyzing subsets based on the expression of CD14 and CD16, we found significantly decreased levels of C.monos and NC.monos and an increased level of I.monos in HS (**Figure 2E**). Taken together, these data highlight previously unappreciated immune cell dysregulation in the peripheral immune system of patients with HS.

### Identification of monocyte subpopulations with elevated CD38 and skin-homing chemokine receptor (shCCR) expression

Machine-learning algorithms can be applied to high-dimensional single-cell data to perform unbiased clustering and statistical comparison of clusters between conditions and to identify previously undescribed subpopulations of cells ^26^. We employed the L1-penalized regression implementation of the cluster identification, characterization, and regression (CITRUS) analysis to confirm our findings from bivariate gating and uncover novel disease-associated immune cell subpopulations in HS. Unbiased clustering of CD45^+^ cells found 11 differentially abundant clusters that corresponded to 9 subpopulations (**Figure 3A & B**). We mapped cells back into t-SNE space and analyzed the expression pattern of lineage markers to determine their identity (**Figure 3C-D; S2**). CITRUS annotations are shown in **Figure 3D** and **Figure S2.** CITRUS results confirmed some results from bivariate gating. Notably, two subpopulations of I.monos were more abundant in patients with HS; and NC.monos, C.monos, and NK cells were decreased in patients with HS (**Figure 3D; S2**).

**Fig.3.**
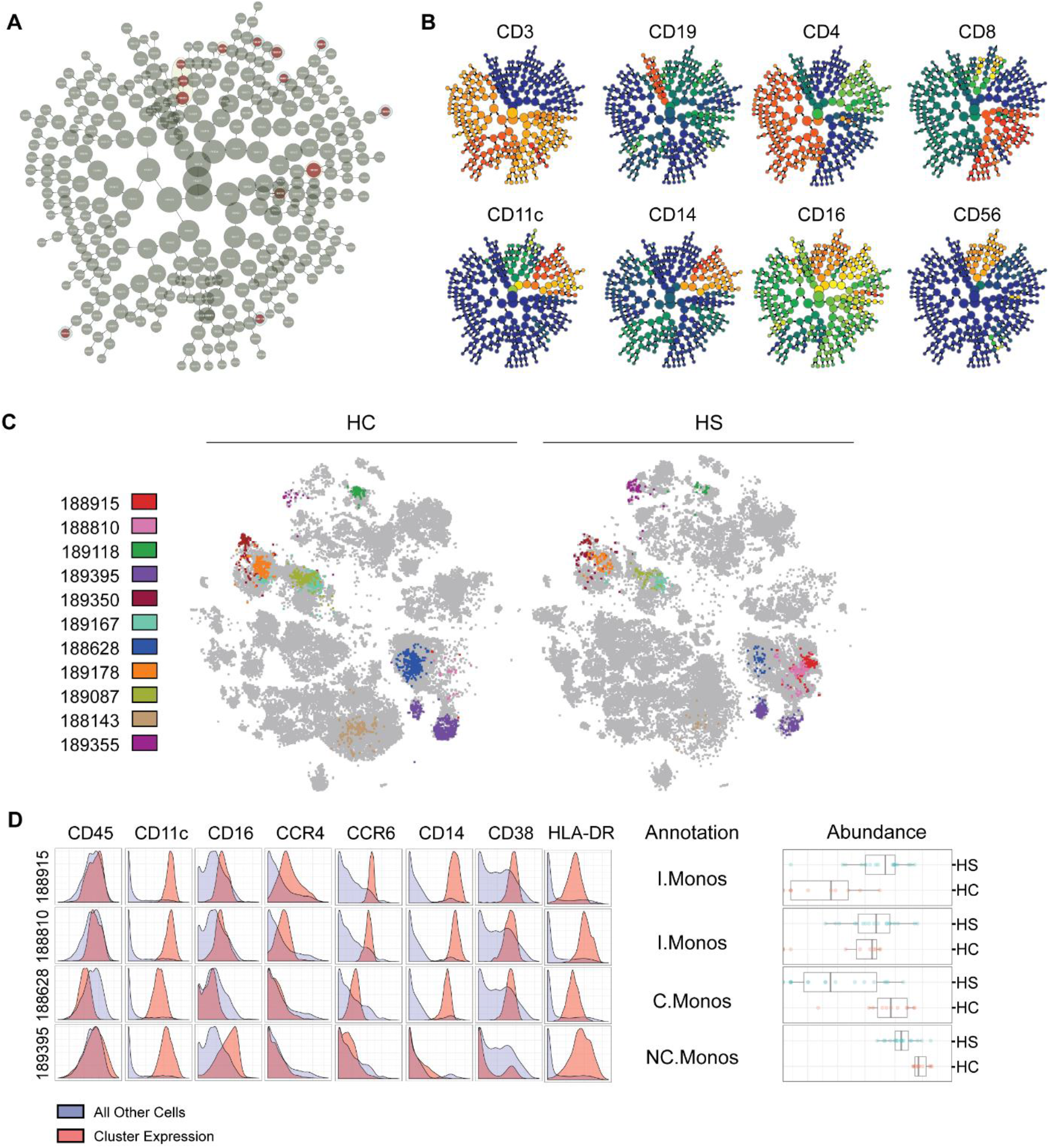
Identification of significantly dysregulated subpopulations in the peripheral immunomes of patients with hidradenitis suppurativa: (A) CITRUS clusters from L1-penalized regression LASSO analysis, statistically significant clusters are highlighted red.; (B) Protein expression plots for lineage markers overlayed on CITRUS clusters; (C) Cells from significant CITRUS clusters mapped to t-SNE plots. All other cells are colored gray; (D) Marker expression histograms for clusters 188915, 188810, 188628, & 189395. Red curves represent cluster expression level and blue curves are the expression of all other CD45^+^ cells. Annotations were made based on both marker expression and t-SNE localization. Boxplots show the abundances of each cluster in HS and HC. Abbreviations. C.monos, classical monocytes; HC, healthy control; HS, hidradenitis suppurativa; I.monos, intermediate monocytes; NC.monos, nonclassical monocytes.

We further assessed the expression of markers that may indicate dysregulated function in monocyte subpopulations (**Figure 3D**). CD38, also known as cyclic ADP ribose hydrolase, is an enzyme that metabolizes NAD+ and has been shown to be required for optimal activation of monocytes and macrophages in response to toll-like receptor (TLR) stimulation, tissue-trafficking, antigen presentation, and phagocytosis ^27–29^. CD38 is induced on immune cells in inflammatory diseases such as rheumatoid arthritis, systemic lupus erythematous (SLE), and multiple sclerosis ^27, 28, 30^. We found that the CITRUS clusters corresponding to I.mono and C.mono subpopulations expressed high levels of CD38 (**Figure 3D**). C.monos normally express appreciable levels of CD38 ^22, 24^. I.monos, on the other hand, do not express high levels of CD38 unless stimulated, highlighting that these I.monos are indeed in an activated state in HS.

Monocyte trafficking to tissues depends on CCR2, which is expressed by C.monos and not I.monos or NC.monos ^24, 31,32^. Under inflammatory conditions, the skin produces cognate chemokines for CCR4 and CCR6, which instructs immune cells to localize to inflamed skin ^33^. We found that CITRUS I.mono subpopulations expressed high levels of both CCR4 and CCR6, while the subpopulation of HS C.monos expressed increased CCR6 (**Figure 3D**). High expression of shCCRs, was not present on the NC.mono subpopulation in HS.

Taken together, these data show that activated subpopulations of I.monos and C.monos are present in the circulation of patients with HS and express elevated levels of shCCRs.

### Expression of shCCRs and CD38 across circulating immune cell subsets in patients with HS

We next wanted to determine whether upregulation of CCR6, CCR4, and CD38 was restricted to specific subpopulations of monocytes identified by CITRUS, or if these changes were reflective of perturbed cell surface expression of the whole monocyte subset population in patients with HS. We also sought to identify other immune cell subsets that have perturbed expression of shCCRs and CD38 that were not identified by CITRUS.

Neutrophils, I.monos and, C.monos from patients with HS had elevated CCR6 (**Figure 4A**). Memory B cells, plasmablasts, I.monos, C.monos, and Th2 cells from patients with HS had elevated CCR4 expression (**Figure 4B**).

**Fig.4.**
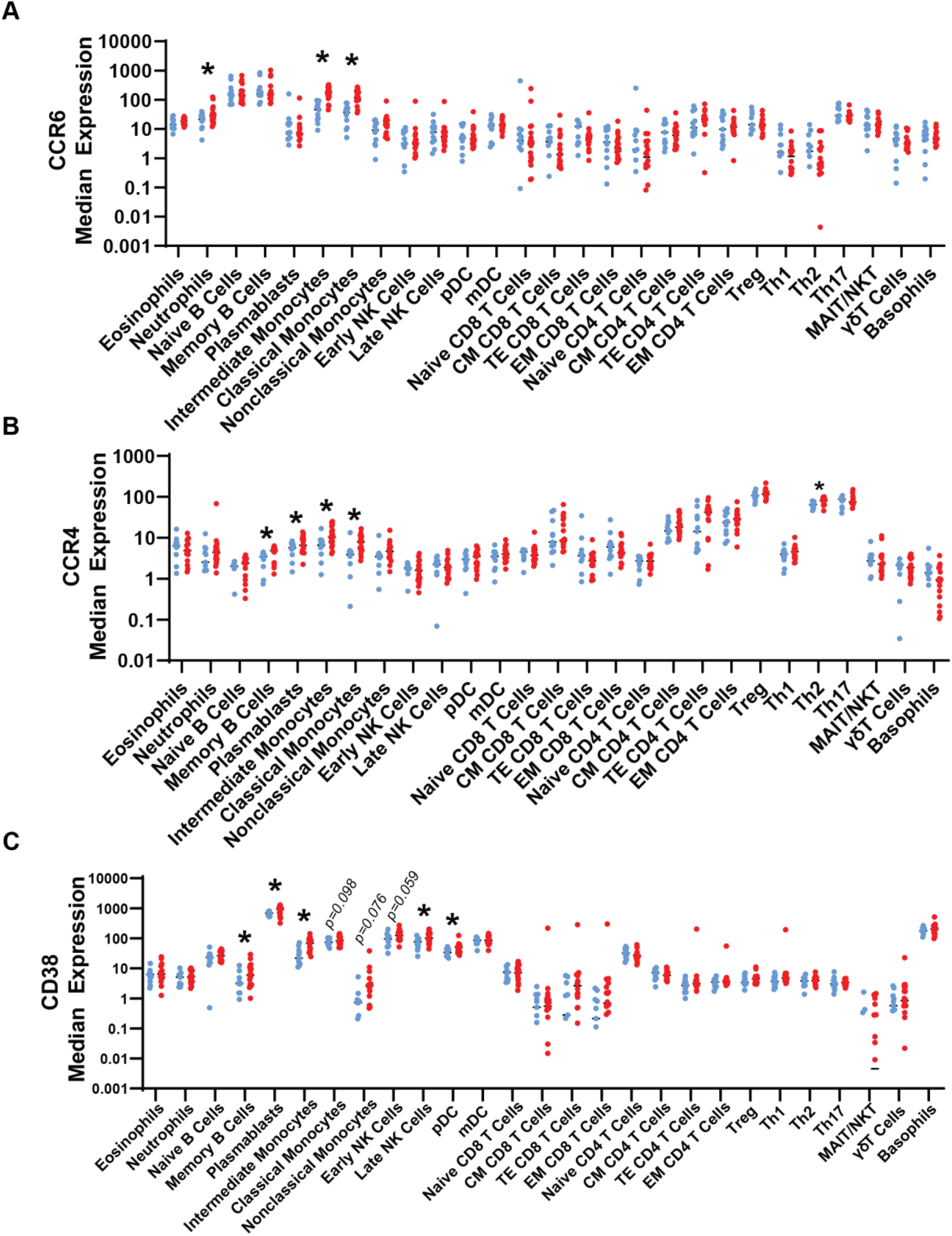
Dysregulated expression of shCCRs and CD38 in immune cell subsets in peripheral blood of patients with hidradenitis suppurativa: Median expression of (A) CCR6, (B) CCR4, and (C) CD38. Asterisks represent *p*<0.05 from unpaired student’s t tests between HC and HS. Abbreviations: HC, healthy control (blue); HS, hidradenitis suppurativa (red).

Memory B cells, plasmablasts, I.monos, late NK cells, and plasmacytoid DCs (pDCs) from patients with HS exhibited increased CD38 expression (**Figure 4C**). CD38 levels of C.monos (*p*=0.098), NC.monos (*p*=0.076), and early NK cells (*p*=0.059) showed a trending increase in HS, approaching statistical significance. (**Figure 4C**). Increased CD38 expression in these cell subsets did not directly correlate with changes in their frequencies, suggesting cell-type specific consequences of elevated CD38, consistent with previous studies of other inflammatory diseases (**Figure 1D, 2E, S1**) ^30, 34, 35^. The increased expression of CCR4, CCR6, and CD38 on I.monos in patients with HS is consistent with our CITRUS findings, and interestingly, it appears that C.monos overall have an increase in CCR4 and CCR6 in patients with HS (**Figure 3;4A-F**).

Taken together, these data highlight enhanced expression of shCCRs and CD38 on multiple circulating immune cell subsets in patients with HS.

### CD38^+^ intermediate monocytes may serve as a circulating biomarker of HS disease progression

We devised a gating strategy to identify CD38^+^ monocytes. Normally, CD38 is not expressed by NC.monos, which we leveraged to define the background level of CD38 in monocyte subsets (**Figure 5A**)^36^. As expected, C.monos from HCs were almost entirely CD38^+^ (**Figure 5A**). We found that the frequency of CD38^+^ I.monos was elevated in HS, which is consistent with our CITRUS results (**Figure 5B**). A discrete population of CD38^+^ NC.monos was present in only some patients with HS (*p*=0.0819).

**Fig.5.**
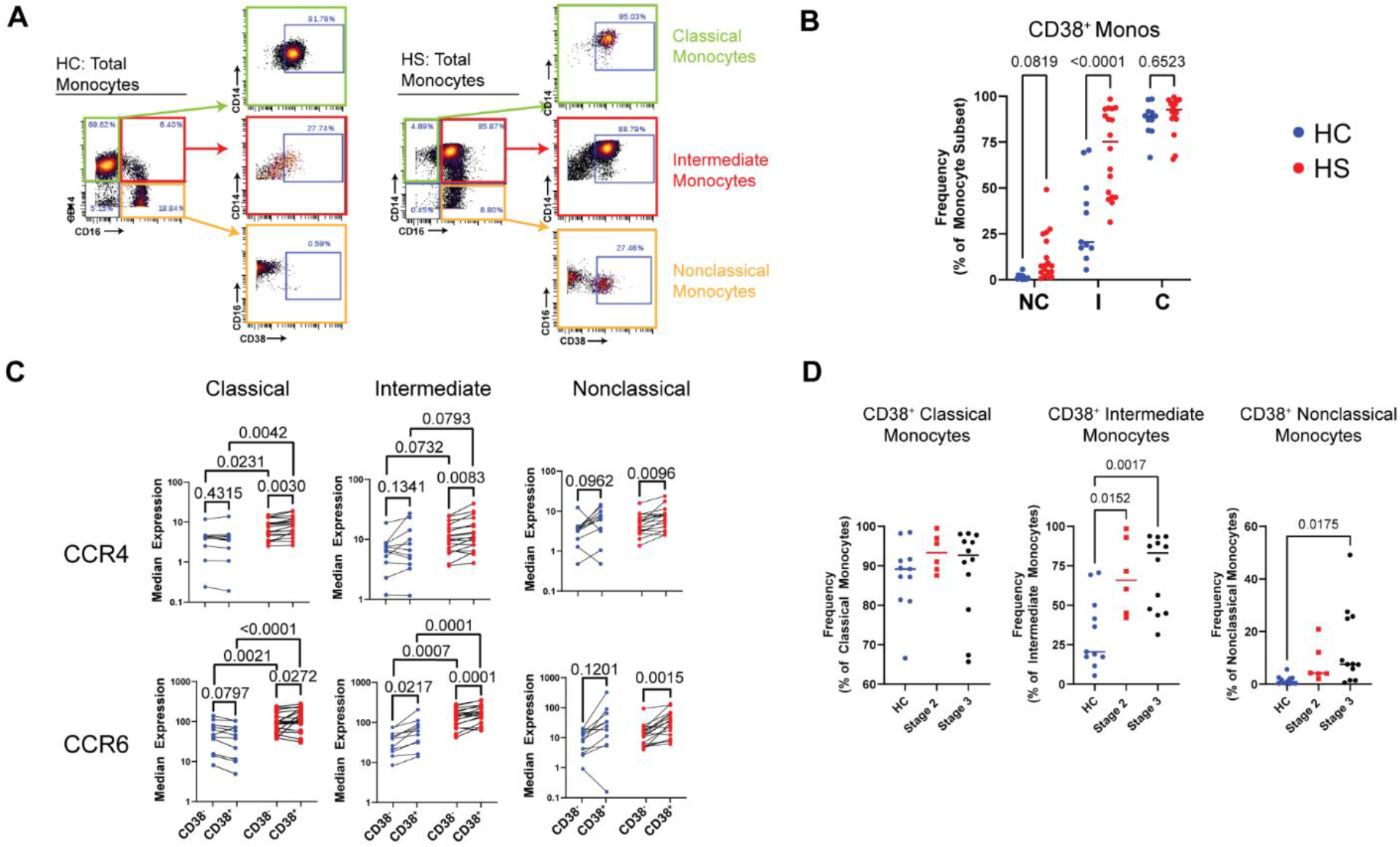
Circulating CD38^+^ intermediate monocytes are elevated in patients with moderate hidradenitis suppurativa, and shCCR expression is increased in monocyte subsets from patients and in CD38^hi^ populations: (A) Gating strategy to identify CD38^+^ monocyte subsets; (B) Frequency of CD38^+^ monocytes as a percent of their respective subset. *P*-values reported are a result of unpaired students t-tests. (C) Expression of CCR4 and CCR6 stratified by HS vs. HC and CD38^hi^ vs. CD38^lo^ in monocyte subsets. *P* values reported are a result of paired students t-tests. (D) Frequency of CD38^+^ monocyte subsets as a percent of their respective monocyte subset stratified by Hurley Stage. *P* values reported are a result of one-way ANOVA with multiple comparisons. Abbreviations: HC, healthy control; HS, hidradenitis suppurativa; NC, nonclassical; I, intermediate; C, classical.

Given that increased CD38 enhances monocyte trafficking and activation ^28, 37^, we wanted to determine if the expression of shCCRs tracked in a disease-associated manner and was associated with CD38 expression (**Figure 5C)**. HS C.monos showed a significant disease-associated increase in CCR4 and CCR6 for both CD38^+^ and CD38^-^ cells. HS I.monos showed a significant disease-associated increase only in CCR6, while increases in CCR4 approached significance (CD38^-^: *p*=0.0732; CD38^+^: *p*=0.0793) (**Figure 5C**). CCR4 and CCR6 expression were elevated in all CD38^+^ monocyte subsets in patients with HS relative to their CD38^-^ counterparts. In HCs, however, only CCR6 expression increased with CD38 in I.monos (**Figure 5C**). Taken together, this suggests that C.monos and I.monos from patients with HS show increased skin-homing capacity that is potentiated in CD38^+^ monocytes.

Current prognostic challenges in clinical management of HS warrant investigations to define biomarkers of disease progression and therapeutic response. To address this gap, we wanted to determine if any immune cell subsets were dysregulated in patients in the earlier Hurley Stage II of HS. Analyzing the frequency of all immune cells identified by our CyTOF assay and CD38^+^ monocyte subsets, we found that CD38^+^ I.monos were the only population that showed a significantly elevated frequency in patients at Hurley stage II (**Figure 5D;S3**). We saw a further increase in the frequency of CD38^+^ I.monos in patients at Hurley Stage III, but this was not significantly different from Hurley Stage II results. Together, these data suggest that CD38^+^ I.monos may serve as an early biomarker of HS disease progression and that C.monos exhibit a disease-associated and CD38-associated increase in skin-homing capacity.

### CD38 is robustly elevated in lesional HS skin, but not perilesional HS skin

We were interested to see if CD38 was elevated in lesional skin from patients with HS. We utilized our recently developed computational resource, HS-OmicsDB (https://shiny.hfhs.org/hsomicsdb/), which aggregates publicly available bulk and single-cell RNA sequencing results from studies of tissues from HCs and patients with HS, to determine if CD38 expression is elevated in HS tissue and what gene signatures are associated with high levels of CD38 in HS (manuscript submitted). We found that CD38 expression was significantly higher in HS lesional (L) tissue than in HS perilesional (PL) and HS nonlesional (NL) skin, as well as tissue from HC skin (**Figure 6A**). We then defined a CD38 gene-signature by finding the genes that significantly correlated with CD38 expression (**Figure 6B)**. A total of 1736 genes had significant positive-correlation with CD38 over all samples (*r>0.5; FDR<0.05*) (**Table S4**). We enriched these significantly correlated genes using MSigDB ^38^, the Human Gene Atlas ^39^, and gene ontology:molecular function (GO:MF) ^40^ databases to understand the gene programs associated with CD38 expression in HS. The genes were enriched for the following MSigDB pathways: allograft rejection, IFNγ signaling, complement activity, IL2/STAT5, and IL6 signaling (**Figure 6C**). Annotation to the human gene atlas found that CD38-correlated genes were enriched for those expressed by CD14^+^ monocytes (C.monos), NK cells, and DCs (**Figure 6D**). These three subsets were decreased in our CyTOF analysis of circulating immune cells in patients with HS, suggesting immune infiltration to HS skin lesions **(Figure 2**).

**Fig.6.**
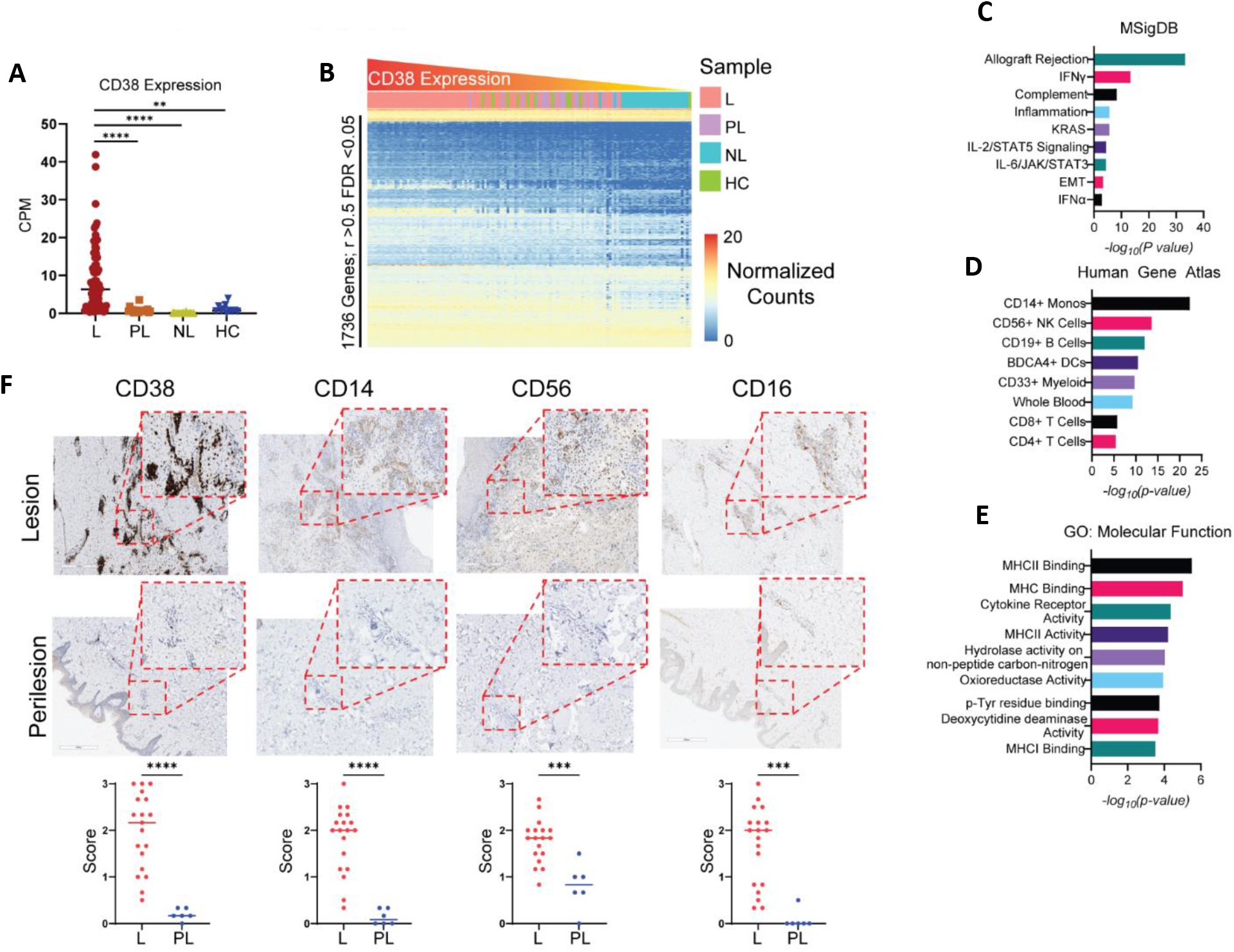
Robust expression of CD38 in lesional HS skin compared to perilesional HS skin: (A) Counts per million (CPM) of CD38 from HS lesional (L), HS perilesional (PL), HS nonlesional (NL), and healthy control (HC) skin. (B) Heatmap of normalized counts of genes significantly correlated with CD38 expression (Bonferroni corrected FDR<0.05). Samples were ordered by CD38 expression from left to right. Significantly correlated genes were enriched using MSigDB (C), the Human Gene Atlas (D), and GO:MF (E). (F) Immunohistochemical staining for CD38, CD14, CD56, and CD16. Staining intensity was scored on a scale of 0 to 3 relative to a positive control and negative control. Each point is the average of three scores from independent raters. Two-way Student’s t test \*\*\*\**p* < 0.0001; \*\*\**p*< 0.001.

CD38 is induced on myeloid cells in response to TLR activation, which enhances cytokine expression and antigen presentation capabilities ^27, 28^. Accordingly, gene GO:MF analysis showed the CD38 gene-signature enriched for the following terms: MHCII, MHCI, and cytokine receptor activity, further indicating that CD38^hi^ lesions are active inflammatory lesions ^4^.

Previous reports have shown increased immunohistochemical (IHC) staining for markers of B cells, DCs, CD8 T cells, and CD4 T cells in lesional HS skin, consistent with our analysis ^41, 42^. To validate the results from our meta-analysis, we performed IHC analysis of CD38 and monocyte (CD14 & CD16) and NK cell (CD16 & CD56) markers on L and PL HS tissues. A total of 19 L and 6 PL samples were stained. As predicted by our meta-analysis, CD38 and monocyte and NK markers were dramatically higher in the L-skin than in the PL-skin (**Figure 6F**). Interestingly, the staining pattern of CD38 suggested that immune and stromal cells may be expressing CD38 in HS L-skin. These results support our meta-analysis and provide compelling evidence that the observed decrease in circulating C.monos, NK cells, and DCs is associated with immune infiltration into HS L-skin.

### Lesional endothelial cells express CD38 and C.monos localize to lesion-specific endothelial cell microenvironments

Normally, C.monos exhibit high tissue trafficking capability due to the expression of CCR2, whereas I.monos and NC.monos exhibit lower tissue trafficking ability ^22, 24, 31^. Since we found that both C.monos and I.monos in HS had increased shCCRs (**Figure 4; 6C**), we wanted to determine which of these monocyte subsets trafficked to the skin in patients with HS. In addition, we wanted to identify which cells expressed CD38 in HS lesions.

To answer these questions, we performed imaging mass cytometry (IMC) with a panel of 31 antibodies (**Figure S4A-B**). We captured 16 regions of interest (ROIs) from 3 matched L and PL tissues. We acquired images for dermal, epidermal (which includes the adjacent underlying dermis), and tunnel regions (**Figure S4C**). We identified 17 clusters of cells annotated into 10 cell types (**Figure S4D, 7A**). Only one cluster expressed requisite monocyte markers (CD11b^+^ CD68^-^ HLA-DR^+^ CD11c^-^) and showed expression of both CD14 and CD16 (**Figure 7A**). To determine if this staining pattern indicated I.monos present in HS skin, we looked for co-staining of CD16, CD14 and HLA-DR. Little to no co-staining was observed, meaning that CD16^+^ cells in HS skin were likely not I.monos (**Figure S4E**). We conclude that the mono cluster in HS skin is predominantly C.monos.

**Fig.7.**
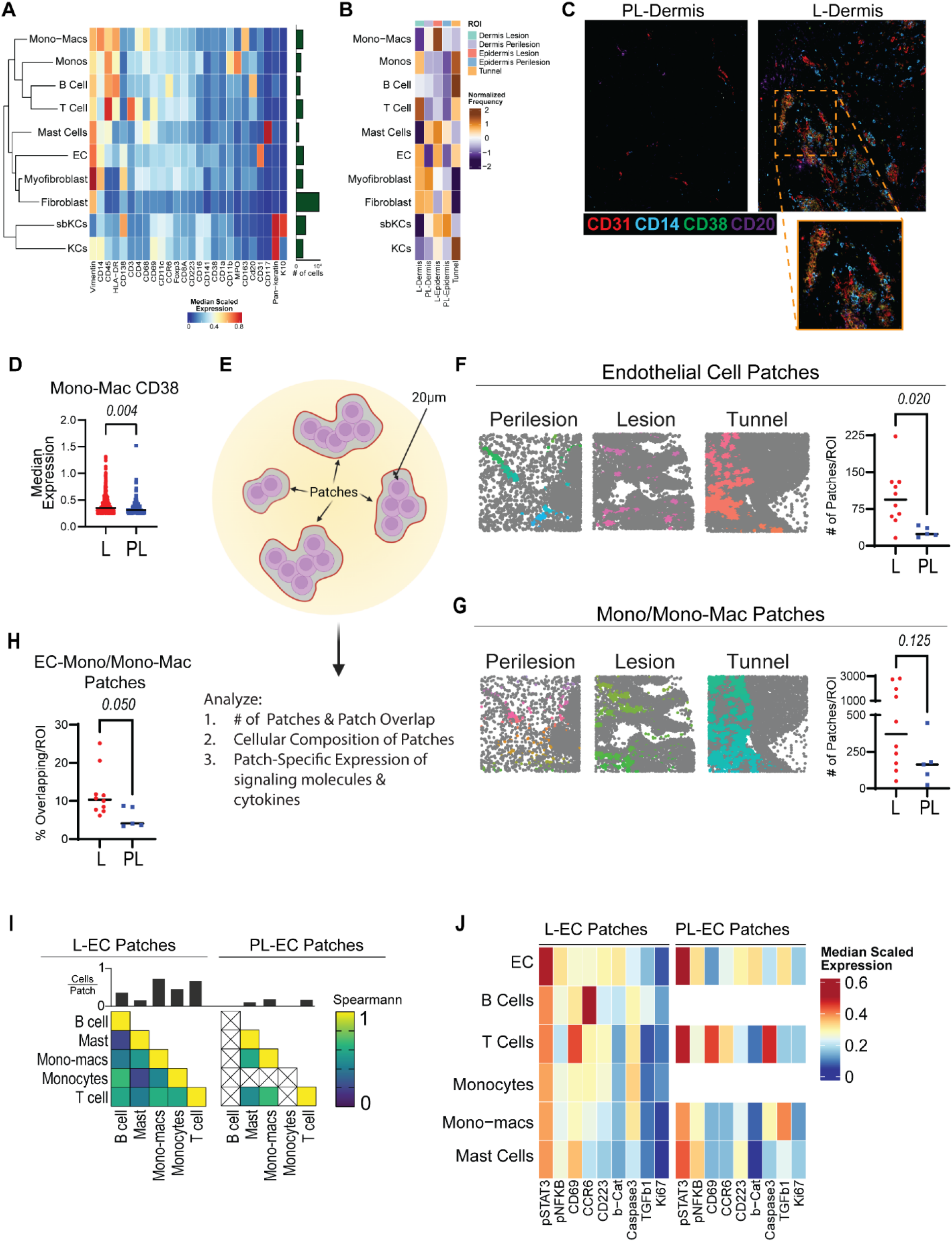
Imaging Mass Cytometry identifies CD38^+^ monocytes, monocyte-derived macrophages, and endothelial cells and dysregulation in the lesional endothelial cell microenvironment in HS skin: (A) Heatmap of marker expression of annotated cell clusters, with bar graph showing the number of cells in each cluster over all captured regions of interest (ROIs). (B) Normalized frequency of each cell type in lesional (L) and perilesional (PL) skin: L-Dermis, PL-dermis, L-epidermis, PL-epidermis, and tunnel ROIs. (C) Representative image showing co-localization of CD31, CD14, and CD38 in PL-Dermis and L-Dermis ROIs. (D) Median expression of CD38 from CD38^+^ mono-macs. (E) Graphical depiction of patch detection method. (F) Left: Voronoi images of endothelial cell (EC) patches colored by unique identification number. Right: Comparison of the number of EC of patches per ROI between L and PL ROIs. *P* value reported is the result of an unpaired students *t* test. (G) Left: Voronoi images of mono/mono-mac patches colored by unique identification number. Right: Comparison of the number of mono/mono-mac of patches per ROI between L and PL ROIs. *P*-value reported is the result of an unpaired students *t* test. (H) Percent of overlapping EC and mono/mono-mac patches in L and PL ROIs. *P* value reported is the result of an unpaired students *t* test. (I) Cellular composition of EC patches. Top: frequency of cells per patch. Bottom: Spearmann correlation of immune cell frequencies in EC patches. Left: L-EC patches. Right: PL-EC patches. (J) Median scaled expression of functional markers on immune cell subsets found in L (left) and PL (right) EC patches. Abbreviations: mono, monocytes; mono-mac, monocyte-derived macrophages.

Comparing the frequencies of immune cells in different ROIs, we found that monos were enriched in L-dermis and L-tunnel regions, monocyte-derived macrophages (mono-macs) were enriched in L-epidermis regions, T cells were enriched in L-dermis and L-tunnel ROIs, and B Cells were most enriched near L-tunnels. Lastly, ECs were enriched in L-dermis, L-epidermis, and L-tunnel regions (**Figure 7B**).

Comparing CD38 staining between L and PL HS tissue, we found a dearth of CD38 expression in PL ROIs and an abundance in L ROIs. CD38 exhibited co-staining with CD31 and CD14 in L-dermal ROIs (**Figure 7C)**. In addition, some B cells in these ROIs also expressed CD38, and we found that CD38 did not co-stain with other markers in our panel (**Figure 7C**). We then quantified the expression of CD38 in monos and mono-macs that expressed CD38 in HS lesions (including L-dermis, L-epidermis, and L-tunnel ROIs) and perilesions (PL-dermis and PL-epidermis ROIs). Using a median expression cutoff of 0.25 to define positive staining, we found that the expression of CD38 in mono-macs was significantly higher in L than in PL tissue (**Figure 7D**). Notably, only 11 CD38^+^ monos were found in PL tissues (**Figure S4F**). Taken into consideration with our CyTOF data, these findings suggest that CD38^+^ C.monos from the blood are preferentially trafficking to HS L-skin over PL-skin. Together, these data show that CD38 is expressed by monocytes, mono-macs, and ECs more highly in HS L-skin than in PL-skin.

Cell surface CD38 depletes extracellular levels of NAD(P)H, which has pronounced effects on the microenvironment by affecting the availability of metabolic and signaling molecules. ^43, 44^. In addition, cells that express CD38 can influence cell localization and motility through homotypic interactions with other CD38-expressing cells and CD31 on ECs. Due to the high CD38 staining in and around ECs, we hypothesized that the EC microenvironment in HS L-skin may be distinct from PL-EC microenvironments. We employed a patch detection algorithm using ECs and mono/mono-macs as the reference clusters. Since PL skin lacked a significant number of monos, we used combined mono and mono-macs (mono/mono-macs) for patch detection. Briefly, patches define a local microenvironment as the space encompassed by a concave hull drawn around a reference cluster that extends 20 μm from the edge of a reference cell (**Figure 7E**) ^45^. From this, we can characterize the number of patches, the overlap between patches, the composition of cells within patches, and the marker expression of cells within patches (**Figure 7E**).

As expected, the number of EC patches was significantly higher in HS L-skin than in PL-skin (**Figure 7F**). We observed a trend of increasing mono/mono-mac patches in L tissue relative to PL tissue (*p*=0.125) (**Figure 7F**). Comparing the proportion of EC and mono/mono-mac patches that spatially overlapped, we found significantly more overlap in HS L-skin than in PL-skin, suggesting that ECs and mono/mono-macs share a microenvironment in HS L-skin (**Figure 6G**).

We then characterized the immune cells in L and PL EC microenvironments. We found a higher frequency and diversity of immune cells in L-EC microenvironments, the one exception being mast cells, which were present equally (**Figure 7I**). While PL EC microenvironments lacked B cells and monos, the correlation of immune cell frequencies was similar between the cells that were present in both L and PL EC microenvironments (**Figure 7I**).

We also compared the median expression of cytokines, signaling molecules, and functional markers in the immune cells present in L and PL EC microenvironments (**Figure 7J**). Notably, mono-macs in L EC patches expressed higher levels of CCR6 (Box1) and lower levels of TGFβ (Box 2) than PL mono-macs (**Figure 7J**). L-ECs also expressed lower levels of TGFβ (Box 3) (**Figure 7J**). Even mast cells, which had similar frequencies in L and PL EC microenvironments, had differing expression of functional markers (**Figure 7J**). Taken together, these data show that the number of EC patches, immune cell composition, and functional status of immune cells differ between HS L and PL EC microenvironments.

Collectively, our IMC and CyTOF data show that C.monos preferentially infiltrate HS L-skin and co-localize in an EC microenvironment that appears to be distinct from that found in PL-skin.

## Discussion

Previous studies have shown that serum effector molecules, autoantibodies, and markers of tissue destruction are elevated in the circulation of patients with HS, highlighting that systemic inflammation is a feature of this disease ^5, 13, 14, 19, 46, 47^. Notably, successful HS therapy reduces circulating inflammatory molecules ^46^. To date, no study has systematically characterized circulating immune cell subsets in patients with HS. In this study, we utilized CyTOF to characterize circulating blood immunomes from patients with HS and in healthy individuals. Our data (1) highlight previously unappreciated changes in circulating immune cell subsets in HS; (2) identify CD38 as a marker of inflammation induced in select circulating and cutaneous cell types; and (3) show a preferential accumulation of I.monos in the blood and C.monos in the skin of patients with HS.

Previous studies have found elevated IL-17 serum protein levels in patients with HS compared to healthy controls ^15, 19^. However, previous studies assaying Th17 cells in circulation have not found significant differences in their frequency or function ^20^. Our data show that not only are the frequencies of Th17 cells higher in patients with HS, but they also have increased capacity to produce IL-17. One explanation for this discrepancy is that the previous studies mainly assessed patients with Hurley Stage II disease, whereas most of the patients in our study were at Hurley Stage III ^20^. The clinical success of IL-17 blockade in HS highlights the importance of this cytokine as a key mediator of disease; however, we noticed that not all patients exhibited increased Th17 cell frequency and altered function, which may explain why some patients respond better to anti-IL17 treatment than others ^46, 48–50^.

We also showed that patients with HS exhibited lower levels of circulating DCs and NK cells. Data from HSOmicsDB and previous studies of HS skin support the idea that the lower levels of circulating DCs is due to infiltration into HS lesions, but future studies are needed to describe the phenotypic and functional properties of DCs in HS lesions ^15, 23, 25^.

NK cells are a poorly studied lymphocyte in HS. Our CyTOF, meta-analysis, and IHC data provide compelling evidence that NK cells infiltrate HS lesions. NK cells perform antibody-dependent cytotoxicity (ADCC) by binding antibody-coated cells via CD16 and releasing cytolytic molecules ^51,52^. NK cells from patients with HS also had elevated CD38 expression, and CD38 enhances ADCC through interaction with CD16^53^. Also, autoantibodies have recently been identified in patients with HS ^54–56^. This suggests an interesting mechanism, where NK cells with enhanced ADCC capacity may act on autoantibodies to promote inflammation in HS. CD38 also enhances extravasation via interaction with CD31 and CD38 on ECs, which is consistent with our observation that these cells are present at lower levels in the circulation of patients with HS^28^.

Circulating levels of C.monos and NC.monos were also lower in patients with HS, while I.mono levels were higher. Previous studies show I.monos in circulation produce high levels of TNFα and IL-1β ^22, 24, 32^. In HS, I.monos expressed high levels of CD38, indicating an activated status, and these cells were the only subset elevated in patients at Hurley Stage II relative to HCs ^27, 28^. Our bioinformatic, IHC, and IMC assays did not suggest that I.monos were infiltrating the skin in patients with HS, suggesting they may play a major role in driving systemic inflammation. Our results showed that C.monos were abundant in HS lesions, which is consistent with their concomitant decrease in circulation. This highlights that discrete cell types may contribute to cutaneous and systemic inflammation, emphasizing the need for drugs that target dysregulated cell types in both the skin and circulation.

CD38 was expressed more highly on several circulating immune cell subsets, including lesional monos, mono-macs, and ECs. In the skin, CD38^+^ ECs increase in response to skin wounding and promote angiogenesis, which is consistent with our observation that they were present at high levels in lesional HS skin ^57^. Another role of CD38 in ECs is to promote the recruitment of immune cells through homotypic interactions with CD38^28^. Accordingly, we found that EC microenvironments harbored more immune cells in HS lesions than in perilesions that also expressed CD38. Thus, in the context of HS, our data show that activated ECs upregulate CD38 and expand within HS lesions, where they are more able to recruit immune cells.

Our IMC analyses were limited by a small sample size of lesions from surgical specimens, which represent only advanced disease. Whether the changes found here are present in earlier stages of disease is unknown, but it will be interesting to understand how the spatial relationships of immune and stromal cells change during the development and progression of HS ^9, 25^.

HS is often resistant to many biologics, such as adalimumab^3^. Targeting a single cytokine may prove insufficient to treat HS because of the multiple inflammatory pathways that are dysregulated in this disease. Depleting dysregulated immune cells may prove more successful. In fact, a trial in patients with SLE who were resistant to anti-TNFα therapy found that targeting CD38 with daratumumab, which depletes CD38 expressing cells, ameliorates disease ^58^. Our data show that multiple circulating and cutaneous cell types in patients with HS upregulate CD38, which is involved in multiple mechanisms that enhance inflammation ^27, 28, 30, 34, 35, 59^. Our IHC and bioinformatic analysis also found a robust increase of CD38 in lesional, but not perilesional HS skin. Depleting CD38^+^ cells may serve to selectively inhibit immune cells that are dysregulated in actively inflamed regions and in circulation of patients with HS. Interestingly, the cells that upregulate CD38 in SLE and HS appear to be different, suggesting that targeting CD38 could provide disease-specific therapeutic benefits ^34^.

## Methods

### Human subjects

Blood samples were obtained from individuals who provided written informed consent, and samples were deidentified prior to processing under guidelines approved by the Henry Ford Health Institutional Review Board. Patients with hidradenitis suppurativa (HS) were enrolled in Detroit, Michigan during regularly scheduled medical appointments. Patients were staged and diagnosed by an expert dermatologist who focuses on treating patients with HS at the HFHS HS specialty clinic. All patients were staged at Hurley stage II or stage III, and demographic information is provided in Table E1. Healthy control (HC) blood was collected at Henry Ford Health from volunteers with no previous history of skin or autoimmune disease. All blood samples were collected in cell preparation tubes (CPT tubes, BD Biosciences, San Jose, CA) containing sodium heparin and were processed for mass cytometry within 4 h of collection.

### Mass cytometry time-of-flight (CyTOF)

Fresh blood was processed and stained using the Maxpar Direct Immunoprofiling Assay (MDIPA) (Fluidigm or The Longwood Medical Area CyTOF core, Boston, MA). Following processing and staining, samples were stored at −80° C. The frozen stained blood samples were placed at room temperature (RT) for 30-50 min until fully thawed and then incubated with 1× Thaw-Lyse buffer (Smart Tube Inc.) at RT for 10 min. The lysis steps were repeated up to 3 times until the pellet turned white evidencing complete lysis of red blood cells. The cells were then incubated with Maxpar Fix and Perm Buffer (Fluidigm) containing 125 nM Cell-ID Intercalator-Ir solution (Fluidigm) at 4° C overnight or up to 48 h before sample acquisition. Groups of samples (4-8/day) were assessed by Helios mass cytometry (Fluidigm) in 4 independent experiments using a flow rate of 45 μl/min in the presence of EQ Calibration beads (Fluidigm) for normalization. An average of 400,000 ± 50,000 cells (mean ± SEM) from each sample were acquired and analyzed by Helios. Gating was performed on the Cytobank platform (Cytobank, Inc. Santa Clara, CA) and FlowJo 10.5.3 (BD Biosciences).

### CyTOF data processing

FCS files generated by CyTOF were normalized and concatenated, if necessary, using CyTOF Software version 6.7. All CyTOF processed files were also uploaded to the Cytobank, a cloud-based analysis platform. Beads, debris, doublets, and dead cells were manually removed by sequential gating per the *Approach to Bivariate Analysis of Data Acquired Using the Maxpar Direct Immune Profiling Assay* (Fluidigm, Technical Note). The total live cells or CD45^+^CD66b^−^ live singlets were selected for viSNE^60^ analysis or gated manually with multiple cell lineage markers to define immune populations..

### Flow Cytometry from PBMCs

Samples of peripheral blood mononuclear cells (PBMC) were thawed rapidly and transferred with 2 ml R1640 medium containing 10% FBS and 50 U/ml benzonase to a fresh 15 ml tube. Then, 5 ml of R1640 medium containing 10% FBS was added, and samples were pelleted by centrifucation for 10 minutes at 300 rpm. The medium was aspirated, and the pellets were resuspended in 2 ml of RPMI 1640 containing 10% FBS and transferred to a culture plate to be placed at 37° C 5% CO2 for 2 hours to rest. Samples were then centrifuged, and pellets were resuspended in staining buffer (2% FBS, 1X PBS). Samples were incubated with Human TruStain FcX block (Biolegend) for 15 minutes and then were stained with cell surface antibodies for 30 minutes at RT. The following conjugated antibodies were used: CD3 (UCHT1), CD4 (OKT4), CD8 (SK1), CCR4 (LZ91H4), and CCR6 (G034E3), all purchased from Biolegend. Following staining, samples were washed twice with PBS. For intracellular cytokine staining, cells were fixed and permeabilized with IC fixation buffer (Invitrogen) and Permeabilization buffer (Invitrogen) using manufacture’s specifications. Samples were then stained with intracellular markers: IL-17 (BL168), TNFα (Mab11), and IFNγ (4S.B3). Flow cytometry assay was performed using a BD FACSCelesta instrument and data were analyzed with the FlowJo V10.2 software.

### Bulk RNA-sequencing Meta-Analysis

Fastq files were downloaded from the Gene Expression Omnibus for accession numbers GSE151243, GSE154773, and GSE155176, resulting in 130 unique samples. Raw sequence reads were adapter trimmed and quality control assessed. The HISAT2 alignment program was used with default settings to align the paired end reads to Genome Reference Consortium Human Build 38 (CRCh38). Only uniquely mapped reads were used for subsequent analyses. Principal component analysis was used to identify potential batch effects arising from the different data sources. DESeq2 was used to normalize counts and identify differentially expressed genes between sample types.

### Immunohistochemistry

FFPE (formalin-fixed paraffin-embedded) tissues were cut to 4 μm sections, placed on slides and deparaffinized. Slides were dried at 60 °C and then incubated in FLEX Target Retrieval Solution (TRS) at an optimized pH (high or low). Staining was performed for CD38 (Cell Signaling E7Z8C; 1:100, TRS-high pH), CD16 (Cell Signaling D1N9L; 1:500, TRS-low pH), CD56 (Agilent 123C3; TRS-high pH), and CD14 (Cell Signaling D7A22T; 1:250, TRS-low pH) on a Dako Autostainer Link. Slides were washed, dehydrated, mounted, and scanned on a Leica Biosystem Aperio CS2 light microscope.

### Immunohistochemistry Scoring

Three independent raters scored immunohistochemistry staining from 0 to 3, where 0 was a complete absence of staining and 3 was equivalent to a positive control. Each data point represents the average of the 3 raters’ scores.

### Tissue preparation and staining for Imaging Mass Cytometry

All tissue samples had previously been formalin-fixed and paraffin embedded for routine diagnostic pathology at Henry Ford Hospital within the past 5 years. Representative regions of interest (ROI) were selected from lesional and perilesional sections using hematoxylin and eosin staining. Two tunnel regions were captured from lesional sections. Tissue sections were cut at 4 μm. Slides were baked for 1 hour at 60 °C and then dewaxed in xylene x3 for 3 minutes each. Tissue sections were rehydrated in ethanol 100% x3 for 3 minutes each, 95% ethanol x2 for 1 minute each, and water x1 for 1 minute. Slides were fixed in 10% neutral buffered formalin for 30 minutes to help prevent tissue movement. Antigen retrieval was performed using Target Retreival Solution, high pH (Dako) for 30 min at 95 °C in a Decloaking Chamber (Biocare Medical). Slides were allowed to cool and placed in wash buffer. Blocking was performed with Background Sniper (Biocare Medical) at RT for 15 minutes. Samples were incubated overnight in a humidity chamber at 4 °C in the primary antibody master mix diluted in TBS with 1% BSA (Sigma Aldrich). Slides were washed twice in 0.1% TWEEN20 (Sigma Aldrich) in TBS, twice with TBS, then stained with Ir-Intercalator-Ir (Fluidigm) in PBS for 30 minutes at RT. Slides were then washed in water for 5 minutes and allowed to air dry. All samples were simultaneously processed and stained.

### Imaging mass cytometry Data Acquisition

Images were acquired using a Hyperion Imaging System (Fluidigm). The largest square area from each ROI was laser-ablated at 200 Hz and subsequent images were rendered using MCD Viewer software (Fluidigm). A total of 16 ROIs from 3 lesional and 3 perilesional sections from patients with Hurley Stage III HS were assayed.

### Imaging Mass Cytometry Data Analysis

Using the Steinbock Docker^45^ container on Linux, we extracted mass cytometry images and ROI annotation information from the raw mcd files. Cell segmentation was performed using DeepCell (Mesmer)^61^ contained in Steinbock, which utilizes a pretrained deep learning model to distinguish between cell and non-cell regions using spatially resolved protein expression from cell nuclei (DNA1 and DNA2) and cytoplasm/membrane markers (ICSK2, CD45, Keratin, K10, K14, aSMA, CD1a, CD3, CD16, CD68, CD163). The cell segmentation process results in a mask, showing the shape and location of the identified cells, sharing the same xy coordinates as the stained imaging mass cytometry slides. For objects identified in the masks, we then extracted mean pixel intensities for the markers used in the staining panel.

The resultant summary features were then read into R4.2.0 using the imcRtools package ^45^. Initial unsupervised clustering was performed using Rphenograph, taking into consideration the expression of cell-type specific markers. Aberrant clusters, which contained a very small percentage of cells with variable expression of non-biologically relevant markers, were removed from downstream analysis to eliminate staining artifacts. After reclustering the remaining cells, phenotypically similar cell clusters were merged into larger metaclusters to define biologically relevant populations. Spatial patches of cell groupings were identified using the imcRtools “patchDetection” function with endothelial cell and mono-mac populations as a reference point.

## Data Availability

Data related to this article are freely available upon reasonable request. Requests should be addressed to the corresponding author.

## Statistical Analysis

Dimensionality reduction, presentation, and statistical analyses for CyTOF and IMC data were carried out using Prism 9 software (GraphPad) or R (v4.2.1). Data was plotted and graphed as mean ± standard deviation of the mean. Tests for normality were conducted and the appropriate parametric or nonparametric tests were utilized. CITRUS analysis was conducted on the Cytobank webserver utilizing its own built-in statistics. Cluster size, event sampling, and cross validation folds were adjusted to minimize the model-error rate. Significant clusters were designated by a false-discovery rate (FDR) <0.01. Deriving a CD38 associated gene-signature was accomplished by finding all genes with significant correlation to CD38, FDR < 0.05 and r>0.5.

## Study Approval

These procedures and protocols were approved by the Henry Ford Health Institutional Review Board (IRB# 12826).

## Author Contributions

P.D. participated in patient recruitment, in experimental design, performed research, collected, analyzed, and interpreted data, performed statistical analysis, and drafted and revised the manuscript. I.H. participated in patient recruitment, in experimental design, performed research, and revised the manuscript. C.Y. participated in patient recruitment and data collection. I.L. analyzed and performed statistical analysis for high-dimensional imaging data and revised the manuscript. J.T, K.S, and N.K. collected data, and revised the manuscript. A.M., R.H., G.V. and A.H. participated in patient recruitment. J.V. participated in data collection, analysis and interpretation, and manuscript revisions. S.D., M.A., W.L., H.L., D.O., participated in manuscript revision. L.Z. and Q.-S. M. supervised all aspects of the work, acquired funding and resources, participated in data analysis and interpretation, manuscript drafting, and manuscript revisions.

## 6. Abbreviations

HC: healthy control
HS: hidradenitis suppurativa
CM: central memory
DC: dendritic cells
EM: effector memory
MAIT: mucosal-associated invariant T cells
NK: natural killer
NKT: natural killer T cells
TE: Terminal Effector
Treg: T regulatory cells
PL: Perilesion
L: Lesion
CyTOF: Cytomertry by time-of-flight
IMC: imaging mass cytometry

## 8. Acknowledgments

We would like to acknowledge the patients who donated to our study. We recognize that without them this work would be impossible and the privilege it is to be able to serve them in this capacity. We hope that through our continued efforts we will relieve the burden of their disease. We would like to thank Yi Yao for providing advice on analysis of CyTOF data. We would also like to acknowledge the members of the Mi Lab and HFHS department of dermatology for their continued support of our work.

